# Small-molecule control of super-Mendelian inheritance in gene drives

**DOI:** 10.1101/665620

**Authors:** Víctor López Del Amo, Brittany S. Leger, Kurt J. Cox, Shubhroz Gill, Alena L. Bishop, Garrett D. Scanlon, James A. Walker, Valentino M. Gantz, Amit Choudhary

## Abstract

By surpassing the 50% inheritance limit of Mendel’s law of independent assortment, CRISPR-based gene drives have the potential to fight vector-borne diseases or suppress crop pests. However, contemporary gene drives could spread unchecked, posing safety concerns that limit their use in both laboratory and field settings. Current technologies also lack chemical control strategies, which could be applied in the field for dose, spatial and temporal control of gene drives. We describe in *Drosophila* the first gene-drive system controlled by an engineered Cas9 and a synthetic, orally-available small molecule.

**Graphical Abstract.**
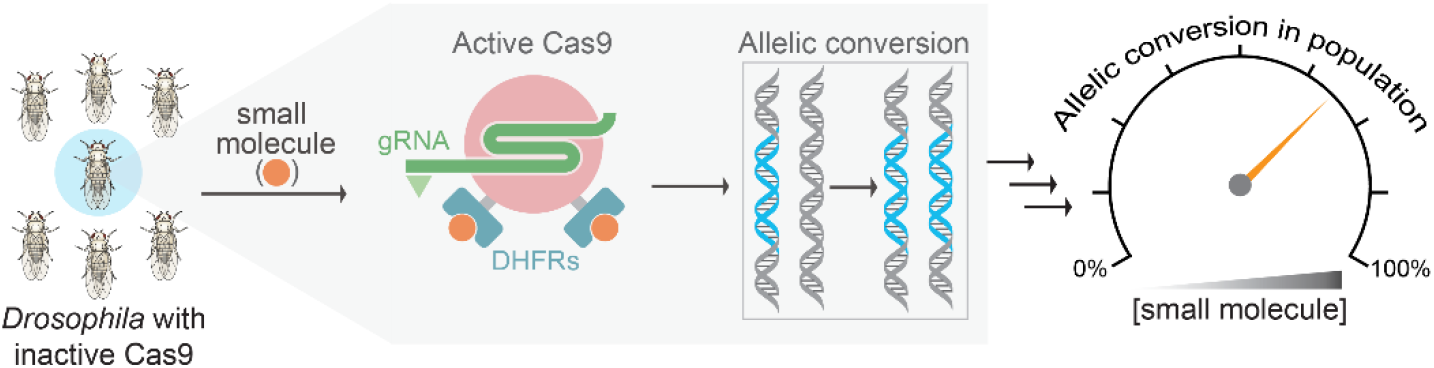

## MAIN TEXT

The Mendelian rules of random assortment dictate that a given gene has a 50% chance of being transmitted to progeny through sexual reproduction (**Fig. 1A**, top), though CRISPR-based gene drives breach this barrier by increasing the inheritance limit towards 100% (i.e., super-Mendelian inheritance). This rate allows rapid gene transmission and is ushering in an era of active genetics^1–3^ (**Fig. 1A**, bottom), making gene drives potentially applicable in everything from basic research^4^ to ecological engineering^4, 5^, including for managing both insect-borne diseases^6, 7^ and invasive pest species^5^ as well as in ecosystem restoration^4, 5^. For example, gene drives have allowed ~100% transmission of antimalarial^6^ or infertility^7^ genes within *Anopheles* mosquito populations in the laboratory, enabling efficient population modification or suppression, respectively. However, multiple concerns and challenges surround the use of gene drives in both laboratory and ecological settings^8^. The unknown consequences of organisms containing engineered gene drives escaping from either the laboratory or their intended ecological residence have prompted intense interest in strategies for spatially containing these organisms^8^. Furthermore, there are no currently existing methods to precisely control the timing of gene-drive activation, to prevent drives occurring in escaped organisms, and to fine-tune the inheritance probability to a value between 50% and 100%. Such fine-tuning of the inheritance probability would allow for both spatial control of the gene drive in the field as well as fundamental studies on the strengths and limitations of gene drives in laboratory settings. For example, an experimental system could be designed to assess the outcomes of gene drives working at different efficiencies in contained cage trials through the use of fine-tuning, and subsequent computational modeling could identify the key optimization parameters.

**Figure 1.**
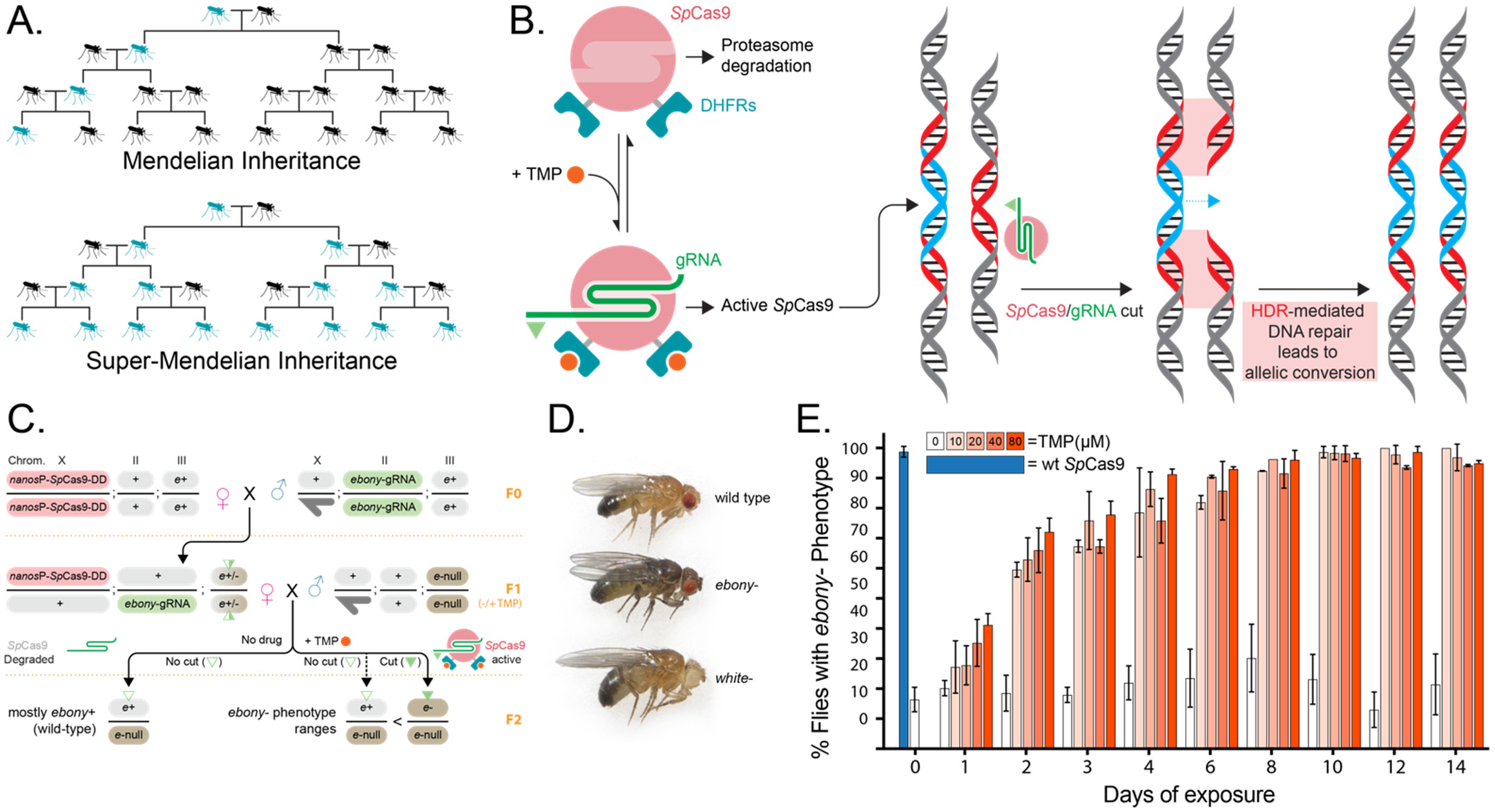
Chemical control of *Sp*Cas9 in Drosophila. **(A)** Super-Mendelian inheritance allows a given genetic trait to propagate exponentially in future lineages. **(B)** A destabilized-domain (DD) system allows small-molecule based dosage and temporal control of *Sp*Cas9 and subsequent gene drives (DD-*Sp*Cas9). In the absence of TMP, DD-*Sp*Cas9 is degraded by the proteasome, while in the presence of TMP, DD-*Sp*Cas9 is active and can induce double-stranded DNA breaks. Repair of the induced cut with the gene-drive-containing template insures the super-Mendelian transmission of the gene-drive construct to the offspring. **(C)** Experimental outline for TMP-activation of *DD-Sp*Cas9 transgenes. F0: Females bearing *nanos*-DD-*Sp*Cas9 transgenes were crossed with males bearing *U6-gRNA* guides targeting *ebony (e).* F1: Female progeny with *nanos-DD-Sp*Cas9 and pFP545 *U6-gRNA* were selected and crossed to *e-/e-* males and fed on food either in the absence (left) or presence (right) of TMP. F2: progeny were scored visually for mutations in *ebony* that indicated activation of DD-*Sp*Cas9 in the germline. A dark-gray half arrow indicates the male Y chromosome. **(D)** Phenotypes of wildtype fly (top), *ebony* mutant (middle), and *white* mutant (bottom). **(E)** Dose-dependent TMP-activation of *DD2-Sp*Cas9 transgenes with *ebony* gRNA. Four days after crossing, flies were transferred to vials of food containing the respective concentration of TMP and were subsequently changed onto fresh food with TMP each day. Offspring were scored for the *ebony* phenotype on the indicated day; *nos*-*Sp*Cas9*(WT)* shown for comparison. Starting on day 2, all values were significant to *p*<0.0001 relative to 0 μM TMP per day of exposure, as determined through a two-way ANOVA with Sidak’s multiple comparisons tests.

CRISPR-based gene drives can achieve super-Mendelian inheritance through a Cas9-induced double-strand break on the wildtype allele that is repaired by copying from the intact gene-drive allele *via* homology-directed repair (HDR), replacing a wildtype allele with the engineered gene (**Fig. 1B**)^1–3^. Therefore, controlling the activity of the initial Cas9-induced DNA break could start or stop the drive process at will. We hypothesized that a gene drive employing a synthetic small molecule that could control Cas9 activity would afford precision control of gene drives for multiple reasons. First, small molecules can provide dosage control of Cas9 activity, and their use in regulating a gene drive would convert the output from singular (i.e., fully-on) to analog, wherein the inheritance probability in the population could be fine-tuned to any value between the off-state (50% inheritance) and the fully-on state (~100% inheritance). Second, the effect of small molecules on the on/off state of Cas9 would be rapid, allowing for precise temporal control of the gene drive by controlling the time the small molecule is made available to the organism. For gene drives meant to propagate lethal or sterile traits to the progeny, temporarily switching off Cas9 could be useful for animal husbandry and population expansion during large-scale studies. Restricting Cas9 activity to a narrow temporal window is also important for proper functioning of the gene drive, as persistent Cas9 activity during development gives rise to drive-resistance alleles or mosaic phenotypes formed by non-homologous end-joining (NHEJ) DNA repair processes^6, 9–11^. Third, by requiring gene-drive inheritance to be contingent on the presence of a synthetic small molecule (as opposed to a naturally occurring small molecule), gene drives can be more easily contained. Finally, small molecule-based methods are already widely used to control mosquito populations in the developing world, in part because of their high efficacy and low production and administration costs. Since mosquitoes regularly frequent human habitats, many insecticides (e.g., pyrethroids) are administered as mosquito net coatings or as aerosols in homes instead of the mosquito’s natural habitat^12^. Thus, small-molecule control methods for gene drives have a strong foundational basis in field settings.

We envisioned to build a small-molecule system for controlling gene drives with the following attributes. First, since gene drives are being implemented on various organisms from insects to mammals^13^, the controller should be adaptable to a wide-range of body temperatures (e.g., from 25°C in insects to 37°C in mammals) and cellular physiologies. Second, the small molecule should be synthetic and not naturally occurring for efficient containment. Third, turning one of the gene-drive components (e.g., Cas9) into a controller itself does not additionally complicate the gene-drive circuitry, potentially lowering chances of failure in field applications. Furthermore, this strategy avoids the use of transcription-based controllers that rely on complex regulation by additional genetic components (e.g. UAS/Gal4 or TetON system). We previously described in mammalian cells a small-molecule-controlled *Sp*Cas9 system built by fusing the structurally unstable *E. coli* dihydrofolate reductase (DHFR) protein to *Sp*Cas9 (DHFR-*Sp*Cas9-DHFR, or DD-*Sp*Cas9)^14^. Upon expression, the DD-*Sp*Cas9 fusion protein is targeted for proteasomal degradation unless the DHFR-binding small molecule, trimethoprim (TMP), is added. Because destabilized domains are efficacious in multiple organisms^15–18^, regulated by a synthetic molecule, provide superior control of DNA-binding proteins over transcription-based controllers^19^, and can control SpCas9 in mammalian cells^14^, we reasoned that this system would provide an ideal gene-drive controller. Furthermore, as the small molecule is required to stabilize DD-*Sp*Cas9 in this default-off system, there is reversible dosage control of *Sp*Cas9 nuclease activity and an added level of safety in case of escaped organisms (**Fig. 1B**).

We developed a DD-*Sp*Cas9 system for *Drosophila* by testing previously-validated clones of DHFR for those that work at lower temperatures^15^ and optimizing them to our system, testing the effective *in vivo* dose ranges of TMP upon ingestion. We explored several reported clones of DHFR and generated transgenic flies with *nos*-DD-*Sp*Cas9 constructs containing 2–3 DHFR domains at varied positions: N-terminal, C-terminal, and an internal loop previously described to tolerate a small protein-domain insertion^20^ (DD1-*Sp*Cas9 through DD4-*Sp*Cas9, **Supplementary Fig. S1)**. We calibrated TMP-dependent *Sp*Cas9 activation by targeting an easily identifiable dark body phenotype that is produced upon mutation of the recessive *ebony* gene. In these optimization experiments, female flies bearing *DD-Sp*Cas9 (DD1-DD4) were crossed to males bearing a gRNA targeting *ebony* under the control of the ubiquitous *U6* promoter (**Fig. 1C**). Subsequently, female progeny carrying both *DD-Sp*Cas9 and *U6-gRNA* transgenes were fed different doses of TMP and crossed to *ebony-/ebony-* males (**Fig. 1C**). The resulting F2 progeny were scored using the visual phenotyping assay for *Sp*Cas9-mediated editing of the *ebony* gene (**Fig. 1D**). While we observed little to no *Sp*Cas9 activity with DHFR fusions for DD1-, DD3-, and DD4-*Sp*Cas9 based on *ebony* mutation rates (**Supplementary Fig. S2**), we observed a greater TMP activation for DD2-*Sp*Cas9, which increases gradually over the first four days and then plateaus at the level of constitutively expressed wildtype *Sp*Cas9 by days 6-10 at all TMP concentrations tested (**Fig. 1E**, **Supplementary Fig. S2**).

We next sought to establish precision control of gene drives using TMP and a modified gRNA-only drive construct called a CopyCat^21^ element that behaves similarly to a gene-drive construct in the presence of a transgenic source of *Sp*Cas9, which is itself transmitted in a Mendelian fashion^21,18^. Briefly, a CopyCat element is composed of a gRNA-expressing gene inserted at the location where such gRNA targets the genome for cleavage; when combined with a transgenic source of Cas9, a Copycat element is capable of copying itself onto the opposing chromosome *via* HDR. CopyCat elements are unable to spread exponentially in a population and increase only additively each generation based on the initial allele frequency of the Cas9 transgene^8^. The small-molecule-controlled *Drosophila* system developed herein consisted of two components: 1) a transgenic source of DsRed-marked *Sp*Cas9 (or DD2-*Sp*Cas9) driven by the *vasa* germline promoter, which was inserted into the *yellow* gene coding sequence; and 2) the GFP-marked CopyCat element containing a gRNA under the control of the *Drosophila U6:3* promoter (**Fig. 2A**). We tested the gene-drive-based inheritance bias of our system by inserting a GFP-marked CopyCat element at the *ebony* and *white* genes (see below), for which homozygous mutants are viable and fertile (**Fig. 1D**). This allowed us to measure inheritance rates by following the fluorescent marker and to evaluate failure events (indels) by scoring the visible phenotype (dark body or white eyes without the GFP marker). First, we crossed males carrying the *Sp*Cas9 (or DD2-*Sp*Cas9) cassette to females containing the *ebony* CopyCat element (F0; **Fig. 2B**). We collected virgin F1 females carrying both the *Sp*Cas9 (or DD2-*Sp*Cas9) and *ebony* gRNA, which were crossed to wildtype males (Oregon-R) for F1 germline transmission assessment using a phenotypic analysis of GFP (CopyCat transgene) in the resulting F2 progeny (**Fig. 2B**; **Supplementary Fig. S3**). This experimental design allowed us to assay the germline inheritance ratios of each single F1 female. Wildtype *Sp*Cas9 showed an ~65% average inheritance for the *ebony* CopyCat element and, as expected, this super-Mendelian inheritance was independent of the presence of TMP (**Fig. 2C**). In contrast, we observed significant TMP dose-dependent super-Mendelian inheritance for the CopyCat element (**Supplementary Table S1**) when combined with the DD2-*Sp*Cas9 construct. In this setup, Mendelian inheritance values of ~50% were seen in the absence of TMP, as expected, and these values reached an average inheritance of ~62% at the maximum TMP concentration (80 μM) (**Fig. 2C**).

**Figure 2.**
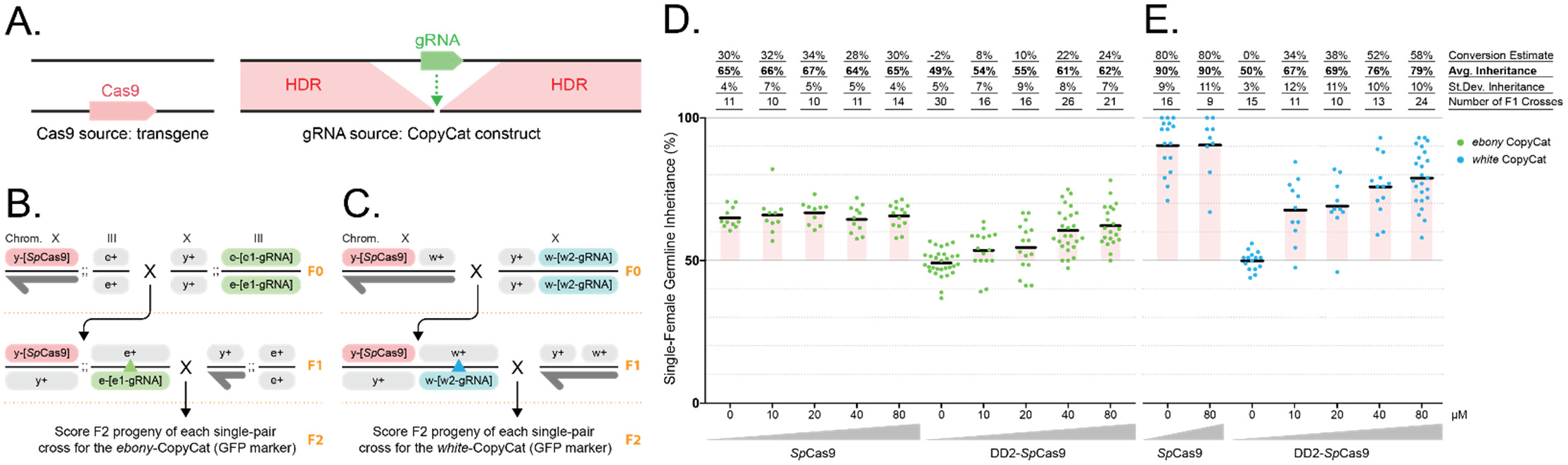
A small-molecule-contingent gene drive. **(A)** Schematic of the CopyCat drive system. The DsRed-marked Cas9 is a static transgene providing Cas9 for mobilizing the GFP-marked CopyCat element by allelic conversion driven by the surrounding homology. **(B,C)** Males expressing *Sp*Cas9 or DD2-*Sp*Cas9 were crossed to virgin females carrying the CopyCat construct. Collected virgin females (Cas9-DsRed + gRNA-GFP) were crossed to wildtype males to score germline transmission rates by screening the GFP marker in the F2 progeny. A dark-gray half arrow indicates the male Y chromosome. **(D,E)** Assessment of gene-drive activity in the germline of F1 females by phenotypically scoring the F2 progeny. Our control line (wildtype *Sp*Cas9), which is not regulated by TMP, displayed super-Mendelian inheritance independent of the presence of TMP. DD2-*Sp*Cas9 showed Mendelian inheritance rates in the absence of TMP (~50%), while TMP treatment triggered an increasing inheritance bias of the gRNA drive element that correlated with TMP concentration for both the *ebony* and *white* CopyCat constructs. Our measurements of inheritance rates allowed us to estimate the allelic conversion rates for our CopyCat constructs, reported on top graph along with the standard deviation (St. Dev.) and the number of F1 crosses performed (n).

To demonstrate that our approach is generalizable to another locus, we generated a second system using a CopyCat construct that targeted the *white* gene, causing lack of pigmentation in the fly eye when disrupted (**Fig. 1D**). We followed a similar experimental approach by crossing males expressing *Sp*Cas9 or DD2-*Sp*Cas9 to females carrying the *white* CopyCat (**Fig. 2D**; **Supplementary Fig. S3**) and raising the progeny on 0 to 80 μM of TMP. The *white* gRNA element driven by *Sp*Cas9 displayed an ~90% average inheritance in both the presence and absence of TMP, reinforcing the conclusion that presence of TMP does not affect *Sp*Cas9 function and suggesting that the components or location of the *white* CopyCat result in a greater copying efficiency than the ~65% of their *ebony* counterparts (**Fig. 2E**). As was the case with the *ebony* construct, DD2-*Sp*Cas9 combined with the *white* CopyCat element had normal Mendelian inheritance rates of ~50% in the absence of TMP (**Fig. 2E**) and, in the presence of TMP, displayed increasing super-Mendelian inheritance rates of 67% (10 μM), 69% (20 μM), 76% (40 μM), and 79% (80 μM), as scored by the GFP phenotype in the F2 progeny (**Fig. 2E**, **and Supplementary Table 1**). This demonstrates that the TMP small molecule provides the desired fine-tuning of the super-Mendelian inheritance rate, which could be used to control gene drive systems based on Cas9. Interestingly, in the absence of TMP, we observed only minimal cutting that was phenotypically evaluated using F2 males of the DD2-*Sp*Cas9 *white* CopyCat experiment (0 μM vs. 80 μM), which was possible because the *white* gene targeted for conversion was located on the same chromosome where the Cas9 was inserted (within the yellow gene coding sequence), and the quantification is given in **Supplementary Fig. S4**. Lastly, we assessed the reversibility of our system by scoring the GFP inheritance of our white CopyCat in a four-generation experiment in which generations F1 and F2 were treated with TMP, while generations F3 and F4 were kept on regular food (**Fig. S5**, **Supplementary Table 2**). Importantly, we observed complete reversibility of our system over subsequent generations. While the TMP-treated F0-F1 displays super-Mendelian inheritance in the F2 progeny, removal of TMP leads to the F4 progeny showing ~50% inheritance of the *white* CopyCat element (**Fig. S5**, **Supplementary Table 2**). To the best of our knowledge, these findings are the first example of gene drives with an analog output wherein the super-Mendelian inheritance can be controlled by the presence of a small molecule.

We herein report the first example of gene-drive elements controlled by a synthetic, orally available small molecule that can regulate the inheritance probability of the genetic element. The controller is transportable to other systems and compatible with current gene drive configurations. The controller activates a gene drive by requiring a deliberate application of a synthetic molecule, providing a new method for increasing the safety of laboratory experimentation^8^ and can be implemented in combination with existing strategies (physical, genetic, ecological)^8^ to provide an additional layer of safety for gene-drive containment. For future field applications, a small-molecule-regulated gene drive could be used to control the spread of gene-drive elements in a circumscribed locale or by vaporizing the drug inside homes as done for mosquito-repellent small molecules. Furthermore, the use of destabilized domains to control Cas9 represents a new tool for spatiotemporal control of Cas9 activity for the *Drosophila* community, and, most importantly, this method should be generalizable to other organisms wherein a *Sp*Cas9-based gene drive has already been demonstrated, such as mosquitoes^6, 7^ and mice^13^. Our first-generation small-molecule-controllers are also amenable to further modifications. For example, while TMP is non-toxic in humans, we observed a developmental delay when flies were fed with TMP, which could be due to toxicity to the insect’s microbiome. Such toxicity could be averted using a prodrug strategy wherein inactive pro-TMP is rendered active by the removal of a “pro-moiety” by an enzyme expressed in the germline of the engineered organism. Multiple orthogonal enzyme-substrate pairs have been reported that are compatible with TMP, including an esterase^22^ and a nitroreductase^23^. Additionally, these first-generation controllers could also be modified to orthogonally control multiple gene drives in a single organism. For example, the controllers could drive two synergistic or antagonistic genes, with the activity of each decided by the dose of the appropriate small molecule. Since our destabilized-domain methodology is extendable to next-generation Cas nucleases (e.g., *Sa*Cas9) and multiple orthogonal destabilized-domain/small molecule pairs are readily available^24–26^, the independent control of two gene drives is now feasible technology.

## ACKNOWLEDGEMENTS

This work was supported by the Burroughs Wellcome Fund (Career Award at the Scientific Interface to A.C.), DARPA (Brdi N66001-17-2-4055), NIH (R21AI126239 to A.C. and DP5OD023098 to V.M.G).

## AUTHOR CONTRIBUTIONS

A.C. conceived the project. B.S.L., K.J.C., S.G., G.S., J.A.W., and A.C. designed and/or optimized the DD-*Sp*Cas9 system in *Drosophila*. V.L.D.A., K.J.C., E.B., V.M.G., and A.C. designed the gene-drive constructs and experiments, which were performed by V.L.D.A., A.L.B., and V.M.G. V.L.D.A., B.S.L., K.J.C., S.G., J.A.W., V.M.G., and A.C. wrote the manuscript, which was edited by all the authors.

## ONLINE METHODS

### Fly rearing and maintenance for phenotype experiments

Flies were raised at 18°C with a 12/12 hour day/night cycle on regular cornmeal molasses medium. Experimental flies were kept at 25°C with a 12/12 hour day/night cycle. For food containing TMP, we used Formula 4-24 Instant Drosophila Food (Carolina Biological Supply Company) reconstituted by adding water or water containing different TMP concentrations (10, 20, 40, 80 μM). Flies were anesthetized to select individuals for crossing and phenotyping and were phenotyped by viewing with a Zeiss Stemi 2000 microscope for gene-editing studies and a Leica M165 FC Stereo microscope with fluorescence for gene-drive experiments. For gene-editing studies as detailed in Figure 1, only flies with the full *ebony* phenotype were scored as *ebony*; all intermediate phenotypes to non-phenotypic flies were scored as wildtype. All these experiments were carried out in a BSL-1 facility at the Massachusetts General Hospital. For the gene-drive experiments as detailed in Figure 2, we used the GFP marker as an indicator of successful conversion. We also tracked the scored mosaic phenotype in the eyes (see Supplementary Table S1). All gene-drive experiments were carried out in a BSL-2 room built for gene-drive purposes at the Biological Sciences Department, University of California San Diego.

### Transgenic line generation and genotyping for phenotype experiments

All injections to generate transgenic flies were performed by BestGene Inc. or Rainbow Transgenic Flies Inc. Transgenic lines for gene-editing experiments (Fig. 1) were generated using site-specific φC31 integration at the ZH2A attB site (2A3) on the X chromosome site using *yw ZH-2A*. For wildtype Cas9 under the *nos* promoter, we used y[1] M{w[+mC]=nos-Cas9.P}ZH-2A w[*] from the Bloomington Stock Center^27^. The *ebony* sgRNA line was pFP545 (ref: ^28^). For gene drive experiments, all constructs were injected into an isogenized Oregon-R (OrR) strain from our laboratory to keep a homogeneous background in all our experiments. All the Cas9 lines were inserted at the same location (*yellow* gene) to ensure comparable Cas9 expression levels. The CopyCat elements flanked by specific homology arms and marked with GFP were inserted in *ebony* and *white* genes. After construct injection, we received the G0 flies in the larval state (80–120 larvae). Once they eclosed, we distributed all G0 adults in different tubes (5–6 females crossed to 5–6 males). Then, G1 progeny were screened for the presence of the specific fluorescent marker in their eyes, which was indicative of marker insertion. Flies positive for the marker were crossed individually to OrR flies (same used for injection) to make a homozygous stock for each transgenic line. Finally, we sequenced the final stocks to confirm the correct integration of the cassette in each line.

### Plasmid construction

The *nanos-DD-SpCas9* transgenic constructs were made using the pnos-Cas9 (Addgene 62208) plasmid. Standard molecular biology techniques were used to build all the constructs analyzed in this work, and sequence information is uploaded on NCBI (accession number: 2183265). Portions of the engineered Cas9 constructs were synthetized using GenScript Inc. DNA Synthesis services.

### Statistical analysis

We used GraphPad Prism 7 to perform all statistical analyses. The two-way ANOVA and Sidak’s multiple comparison tests were used (see Supplementary Table S1 and S2).

