## Supplementary Information for "Small-molecule control of super-Mendelian inheritance in gene drives"

Construct generated for phenotypical assessment of TMP-regulated *Sp*Cas9 (Fig. 1)

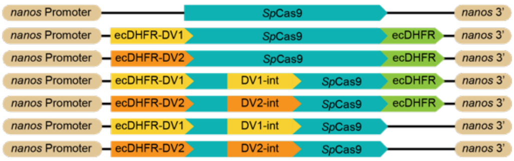


| Construct (informal) | Construct (formal) | Construct Code | Reference |
|---|---|---|---|
| NOS-Cas9 | pNOS Cas9 | NOS-Cas9 (control) | Port et al. 2014 |
| NOS-DV1-recoded | pNOS_N_Clone-10-DHFR_Cas9_C-DHFR | DD1-*Sp*Cas9 | - |
| NOS-DV2-recoded | pNOS_N_Clone-22-DHFR_Cas9_C-DHFR | DD2-*Sp*Cas9 | - |
| NOS-DV1-3Domain | pNOS_N_Clone-10-DHFR_Clone-10-DHFR_Cas9_C-DHFR | DD3-*Sp*Cas9 | - |
| NOS-DV2-3Domain | pNOS_N_Clone-22-DHFR_Clone-22-DHFR_Cas9_C-DHFR | DD4-*Sp*Cas9 | - |
| NOS-DV1-C-term-removed | pNOS_N_Clone-10-DHFR_Cas9 | DD5-*Sp*Cas9 | - |
| NOS-DV2-C-term-removed | pNOS_N_Clone-22-DHFR_Cas9 | DD6-*Sp*Cas9 | - |

Construct generated for gene drive assessment of TMP-regulated *Sp*Cas9 (Fig. 2)

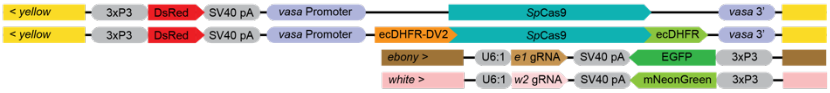


| Construct (informal) | Construct (formal) | Construct Code | Reference |
|---|---|---|---|
| pVG-182 | pVG182_*vasa*-Cas9_in_y1_DsRed | *vasa*-*Sp*Cas9 (control) | - |
| pVG-312 | pVG312_*vasa*-Cas9-DV2_in-y1_HomologyRecoded | DD2-*Sp*Cas9 | - |
| pVMG-51 | pVMG0051_*ebony*-CopyCat-gRNA-e1_GFP | *ebony*-CopyCat | - |
| pVMG-90 | pVMG0090_*white*-CopyCat_gRNA-w2_Dm-mNeonGreen | *white*-CopyCat | - |

**Supplementary Figure S1. Summary of the constructs used in this study**. All constructs contain the same *Sp*Cas9 protein sequence. The *Escherichia coli* dihydrofolate reductase (ecDHFR) domains were added to our wildtype *Sp*Cas9 to generate the DD1-4-*Sp*Cas9 versions that control *Sp*Cas9 expression based on the presence of trimethoprim (TMP). The DD1-4-*Sp*Cas9 were tested based on ebony mutation rates, and the optimal construct, DD2-*Sp*Cas9, was carried forward into the gene drive experiments. For the gene drive experiments (Fig. 2), a red marker (DsRed; red box) driven by an eye promoter (3xP3; grey box) was used for tracking the expression. Both *Sp*Cas9 constructs are flanked by yellow homology arms (yellow boxes), although any allelic conversion process occurs in this transgene. We tested two different CopyCat elements that both carried one gRNA (targeting *ebony* or *white* genes) driven by the U6:1 promoter (grey box) and a green marker (EGFP or mNeon-Green) driven by an eye promoter (3xP3; grey box). Both CopyCat constructs contain homology arms (brown or pink boxes) to allow propagation of the CopyCat element after cutting.

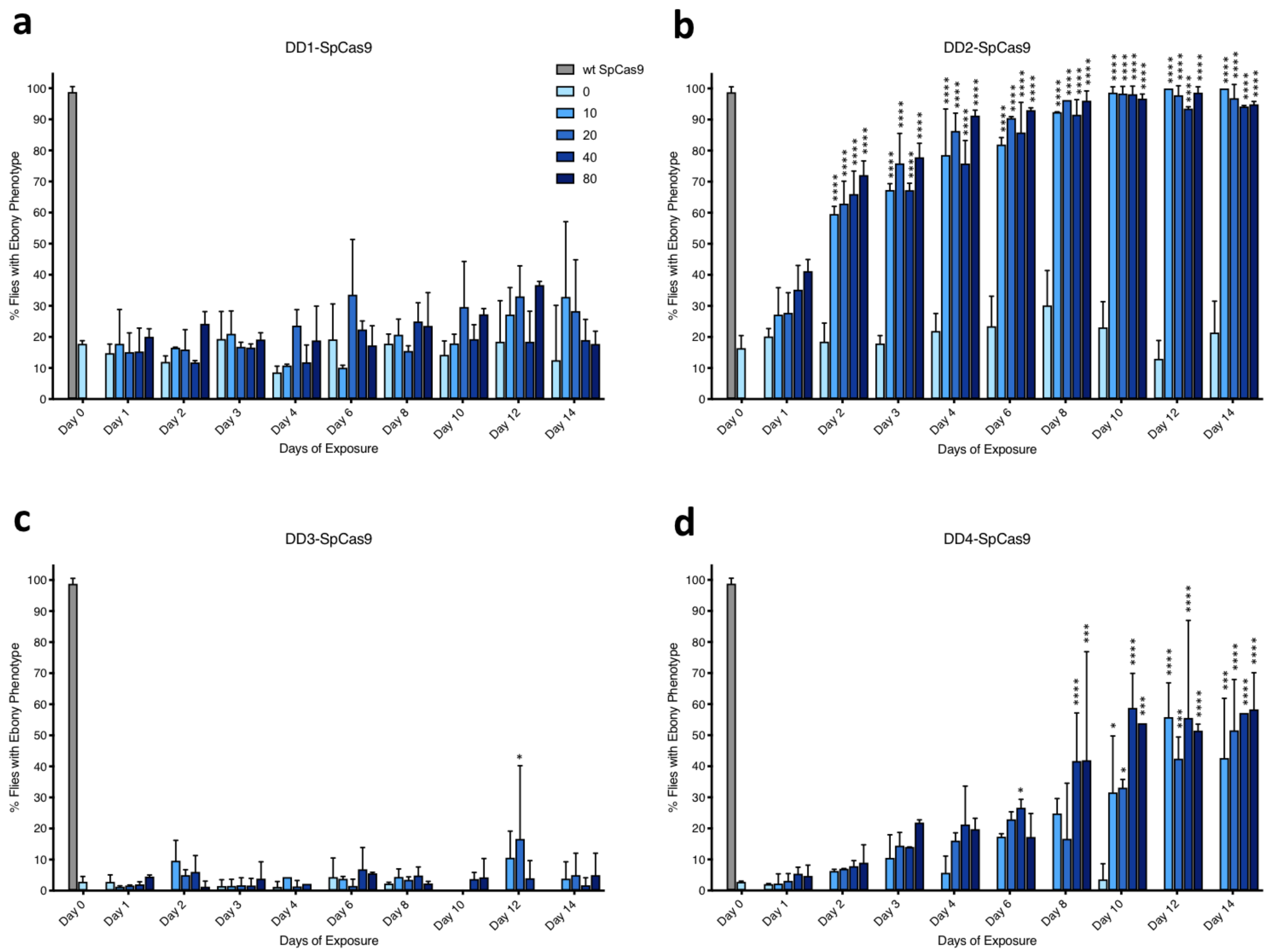


**Supplementary Figure S2. The effect of TMP concentration on the activity of DD1, DD2, DD3, and DD4-*Sp*Cas9 constructs in *Drosophila*.** Dose-dependent TMP activation of all DD-*Sp*Cas9 transgenes with *ebony* gRNA as described in **Figures 1d** and **1e**. Asterisks indicate P-value in comparison to 0 μM TMP per day of exposure (* P<0.05, ** P<0.01, *** P<0.001,**** P<0.0001), determined through two way anova. DD1-*Sp*Cas9 and DD4-*Sp*Cas9 versions displayed low editing activity based on ebony mutant rates, and the DD3-*Sp*Cas9 had unappreciable gene addition. DD2-*Sp*Cas9 showed the highest ebony mutation rates and was chosen as the best candidate for gene drive experiments.

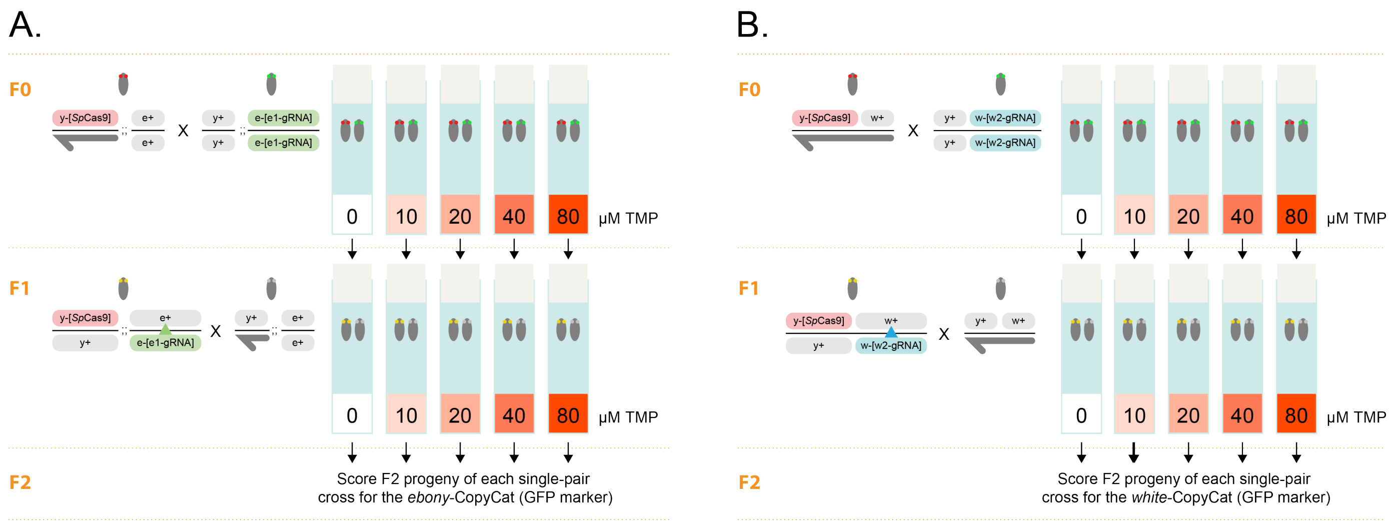


**Supplementary Figure S3. Detailed feeding scheme for dosage control in the CopyCat experiments.** **(A)** *ebony* CopyCat **(B)** *white* CopyCat **(A, B)** F0 crosses between *Sp*Cas9 or DD2-*Sp*Cas9 (DsRed – red eyes) and the CopyCat flies (GFP – green eyes) were kept for four days in vials containing Formula 4-24 Instant Drosophila Food reconstituted with water. The food additionally contained DMSO (control) or the different concentrations of TMP as indicated. After four days, the parents were discarded to collect virgin F1 females carrying both the *Sp*Cas9 (or DD2-*Sp*Cas9) and gRNA elements tracked by DsRed and GFP markers (both constructs – yellow eye in the panel). F1 virgins were single-pair crossed to wildtype males that did not carry any fluorescence (wildtype – grey eyes), using the same DMSO or TMP conditions as the F0 crosses (follow arrows). Lastly, the green fluorescent marker was scored as an assessment of F2 inheritance.

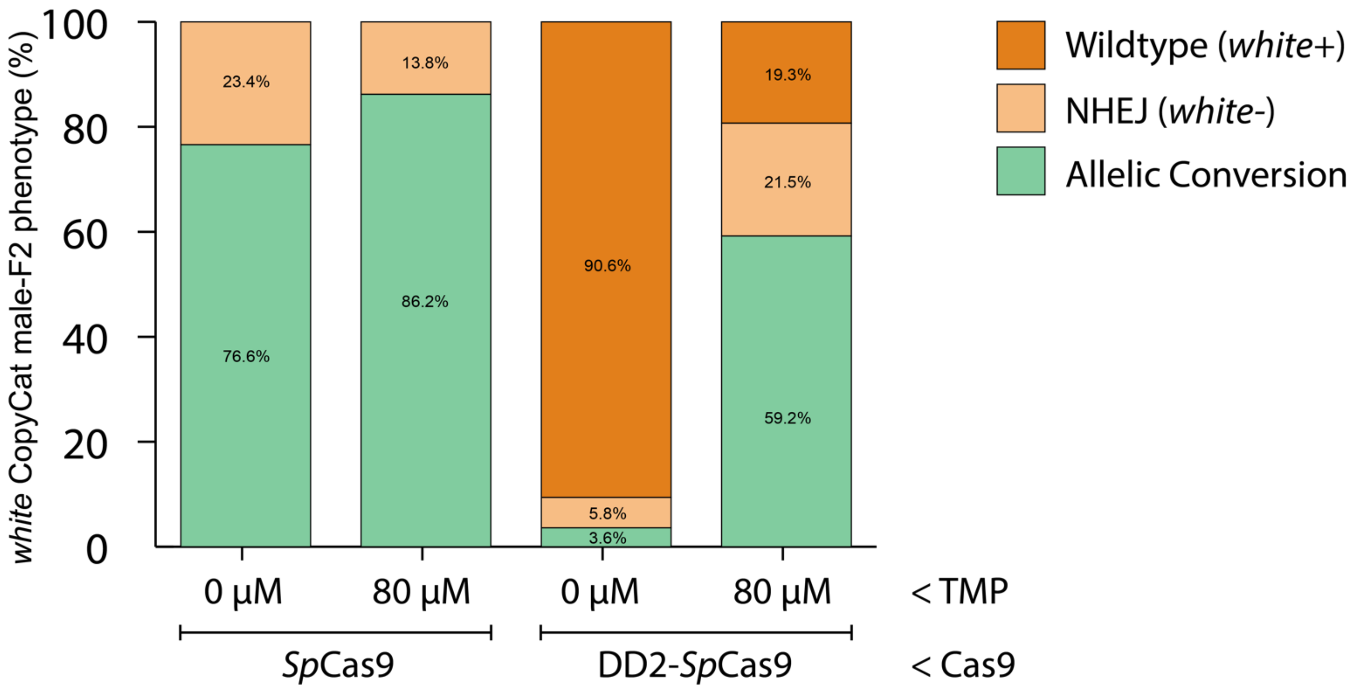


**Supplementary Figure S4. Cutting and conversion efficiency analysis of *Sp*Cas9 and DD2-*Sp*Cas9 in the *white* CopyCat experiment.** Since the *white* gene targeted for conversion is located on the same chromosome where the Cas9 is inserted (within the *yellow* gene coding sequence), we can analyze the receiver chromosome specifically by looking at the F2 males. The results were graphed according to three possible categories based on the phenotypic readouts: 1) Allelic conversion – alleles that were converted (presenting as GFP+ and white eye), 2) NHEJ – alleles that were cut but were not converted (presenting as GFP- and white eye), and 3) Wildtype – these males displayed wildtype eyes (red) and were GFP-, suggesting that these alleles were not acted upon. The cutting efficiency ([conversion + NHEJ] / total) for *Sp*Cas9 was 100% in both conditions (without and with TMP). DD2-*Sp*Cas9 instead displayed about a 9% cutting efficiency without TMP, suggesting a minimal leakiness of the system, and a 79% cutting efficiency in the presence of TMP, which is much lower than the 100% observed for *Sp*Cas9. Considering instead the conversion efficiency (conversion / [conversion + NHEJ]), *Sp*Cas9 showed 77% and 86% for the TMP- and TMP+ conditions, respectively. In contrast, DD2-*Sp*Cas9 showed a much lower 38% conversion efficiency in the absence of TMP than the 73% observed efficiency in the presence of TMP, which is comparable to the values observed for *Sp*Cas9 (see calculation on **Supplementary Table 1)**.

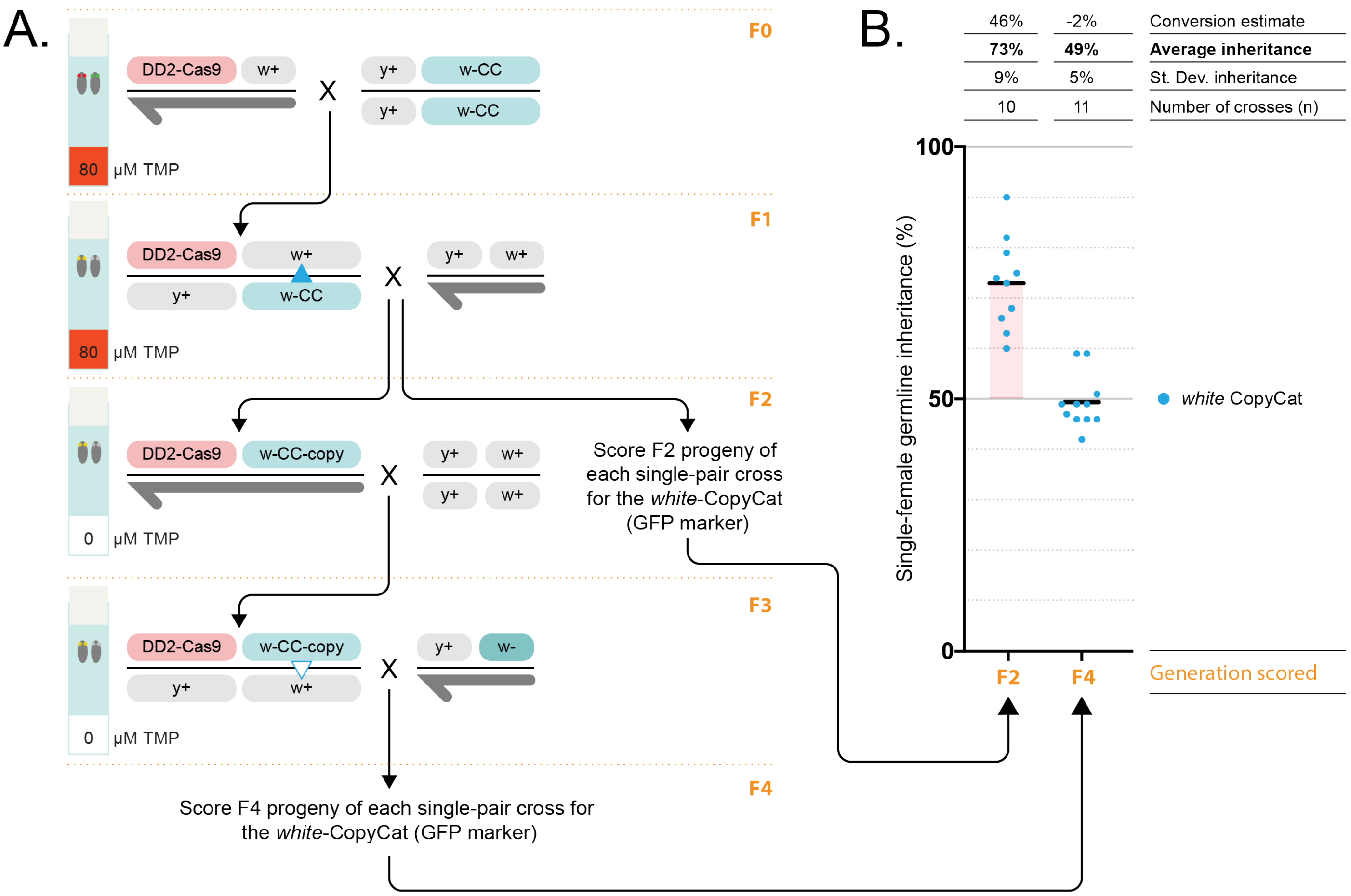


**Supplementary Figure S5. Reversibility of the DD2-SpCas9/TMP-based gene drive system.**

**(A)** Cross scheme to test reversibility in subsequent generations: F0 parents and their F1 progeny are raised in presence of 80 μM TMP, gene drive allelic conversion happens in the germline of the F1 females (solid blue triangle) and is assayed by scoring the frequency of GFP observed in the F2 progeny. F2 males displaying both fluorescence (converted chromosome) were crossed to wild type females in absence of TMP, and resulting F3 females carrying both the DD2-*Sp*Cas9 and the white CopyCat are crossed to *white-* males to score their F4 progeny for inheritance of the GFP marker. **(B)** Inheritance observed in the F2 and F4 generation representative of the gene drive conversion happening in the F1 and F3 female germlines indicated in **(A)** with a solid blue triangle (active conversion) and white triangle (low to no conversion), respectively. Super-Mendelian inheritance is observed in the F2 generation (with TMP) while no conversion and a Mendelian inheritance ratio (~50%) is observed in the F4 progeny (no TMP).
